# Quantifying performance inflation from hidden sample dependence in biomedical image classification benchmarks

**DOI:** 10.64898/2026.10.01.755883

**Authors:** Lukáš Jochymek, Markéta Vašinková, Petr Gajdš

## Abstract

Machine learning benchmarks often treat image files as independent observations, although multiple files may represent repeated views or other related samples. Prior work has shown that non-independent partitioning can inflate model performance, but the sensitivity of a biomedical benchmark to progressively increasing train–test relatedness is less often quantified. Here, we quantified how apparent classification performance changes when recoverable sample groups are progressively allowed to cross training, validation, and test partitions. Using source structure and filenames, we reconstructed analysis groups in a microscopy corpus compiled from multiple sources and generated five class stratified partitioning conditions, ranging from an intact group reference to random assignment of individual files. We applied the same training recipe within each model family to 129,173 images spanning 14 classes of blood cells and hematopoietic cell states. For a pretrained Vision Transformer, random assignment of individual files increased the mean test macro F1 from 0.853 to 0.977 and accuracy from 0.851 to 0.990 relative to the intact group reference. Apparent performance increased progressively across the intermediate conditions. The endpoint pattern also appeared with a ResNet-50 backbone, and the largest apparent gains occurred in immature granulocytic and blast classes. A descriptive analysis within the random assignment condition further showed higher scores among test images that had another member of the same group in the training set, although the exposed and unexposed subsets were not exchangeable. The grouping workflow, exact split definitions, code, and compact result files are released as HSD-Bench to support reproducible evaluation of hidden sample dependence.

## Introduction

Machine learning benchmarks are commonly represented as collections of labeled files, but a file is not always an independent observation. In biomedical imaging, multiple files may correspond to repeated focal planes, acquisition variants, derived crops, image patches, or other representations of the same biological object or acquisition context. If such relationships are not retained as structured metadata, random assignment by file can place related observations across the training and test partitions. Apparent test performance may then partly reflect information associated with an object or acquisition already represented in training rather than generalization to independent observations. Hidden sample dependence is a recognized source of optimistic performance estimates and poor reproducibility in machine learning [1, 2].

Blood cell image classification provides a concrete setting in which this problem is biologically meaningful and experimentally measurable. Many resources in this area are described as white blood cell (WBC) datasets, although they may also contain other blood and hematopoietic cell types or states. A single physical cell may be represented by several focal planes, repeated acquisitions, or derived crops. These alternative views can be useful during training because they expose a model to variation in focus, appearance, and acquisition conditions, but they should not automatically be treated as independent test observations. If one view of a cell is used for training and another view of the same cell is used for testing, the resulting score addresses a different generalization question from performance on previously unseen cells.

Several public blood cell resources illustrate why the unit of independence must be documented explicitly. The Raabin-WBC study notes that the same white blood cell may appear in more than one image [3]. Other datasets contain many cell images acquired from a limited number of patients or smears [4]. Datasets that appear distinct may also share an underlying source; for example, BloodMNIST is derived from the Barcelona peripheral blood cell dataset [5, 6]. These examples do not imply that patient-level separation is required for every morphology task. Instead, they show that the appropriate split unit depends on the intended generalization claim.

For the classification of individual cell morphology, repeated views, derived crops, or acquisition variants of the same physical cell should remain in one partition. Separating patients may also be useful when patient metadata are available, but it addresses a stricter question than generalizing to previously unseen cells. For diagnosis at the patient level, prediction of disease state, or prediction of clinical outcomes, all images from one patient should remain in one partition. Images from the same smear or specimen should also remain together when appropriate. Thus, the split unit determines the type of independence represented by the reported score. Fig 1 summarizes this distinction.

**Fig 1.**
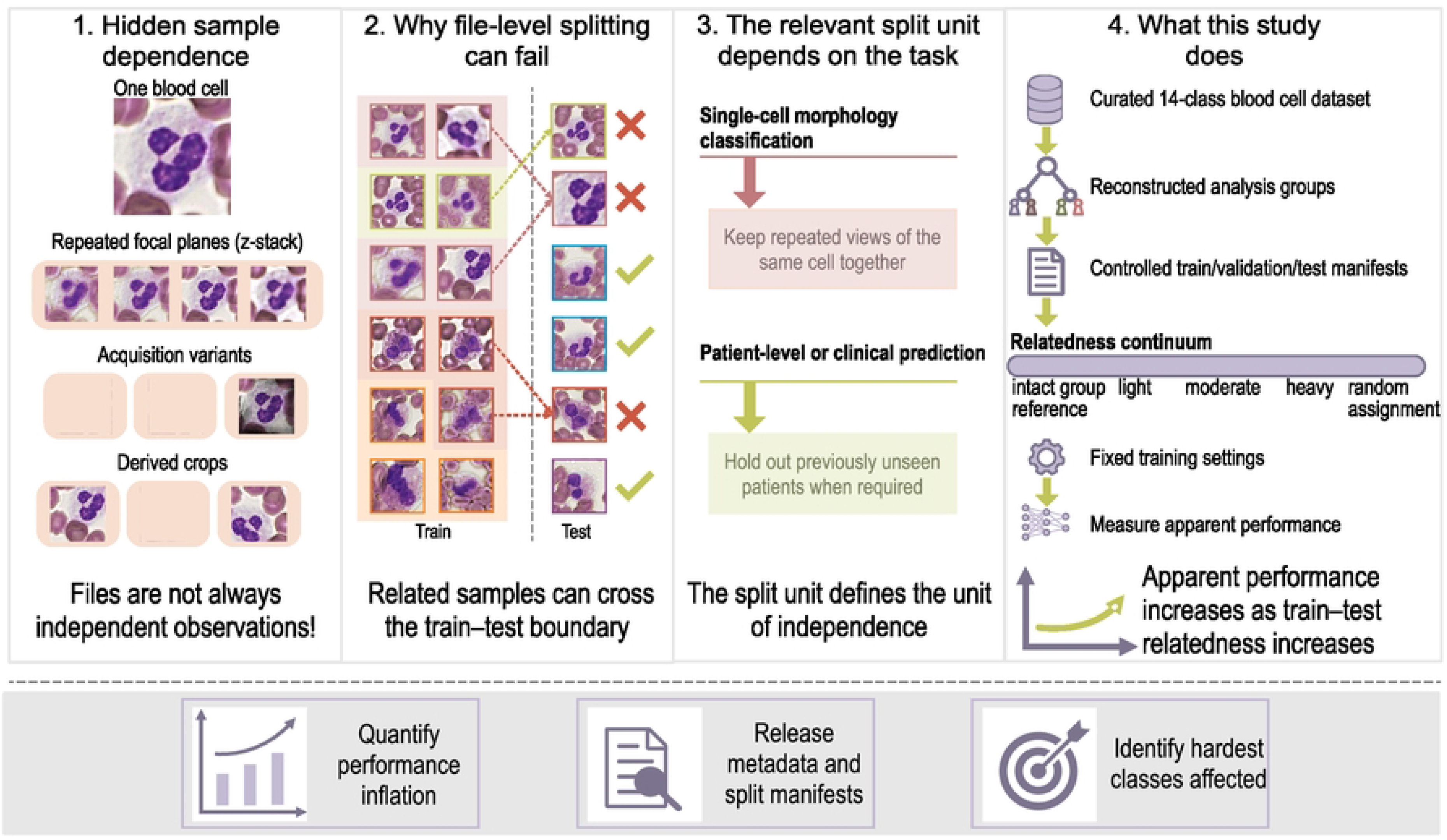
Conceptual overview of hidden sample dependence in biomedical image classification benchmarks. Image files may represent repeated focal planes, acquisition variants, derived views, or other related representations of the same biological object or acquisition context. Under random file-level assignment, members of a related group can occur in both training and test sets. The relevant unit of independence depends on the task: repeated views of the same cell should remain together for single-cell morphology classification, whereas patient-level or clinical prediction requires holding out previously unseen patients. The present study reconstructs recoverable analysis groups, generates controlled train/validation/test partitions, and measures how apparent performance changes as cross-partition relatedness increases. The blood cell microscopy images used in this illustration originate from the publicly available dataset by Park et al. [7].

Despite this distinction, benchmark partitions are often reported only as numerical ratios, such as 70/15/15 or 80/20. These ratios describe how many files were assigned to each partition but not what was held independent. The same ratio can refer to unseen images, cells, acquisitions, smears, or patients. Each choice supports a different generalization claim. Without provenance metadata that describe sample relationships, later users cannot reconstruct partitions that keep related groups together or determine whether related files occurred in both training and test sets. This limits reproducible reuse, complicates comparisons across studies, and makes benchmark scores harder to interpret.

Previous work has established that inappropriate or non-independent partitioning can produce optimistic machine learning estimates [1, 2]. The question addressed here is narrower: how sensitive is one biomedical image benchmark to a graded increase in recoverable sample relatedness when the source corpus and training recipe are otherwise held constant? A graded design can distinguish a systematic relationship from a comparison of two arbitrarily chosen splitting protocols and can reveal whether the largest differences are concentrated in particular biological classes.

The objective of this study was to reconstruct recoverable sample groups in a multisource blood cell microscopy corpus, progressively allow members of these groups to cross dataset partitions, and quantify the resulting changes in apparent classification performance. We generated training, validation, and test partitions ranging from an intact group reference to random assignment of individual files while keeping the training procedure and model settings fixed. We then measured how apparent classifier performance changed as the relatedness between the training and test data increased. We also examined whether the observed pattern was consistent across transformer and convolutional backbones and across individual cell classes. We release the workflow as HSD-Bench, an open-source tool for reconstructing recoverable sample relationships, generating controlled partitions, and quantifying how partitioning choices affect apparent benchmark

performance.

## Materials and methods

### Data description

This study used a curated multi-source microscopy corpus containing 129,173 images in 14 classes representing blood cells and hematopoietic cell states. The corpus was originally assembled for the development and evaluation of biologically informed classification models and combines three public sources: AML-Cytomorphology [4, 8], Raabin-WBC [3, 9], and the multi-focus WBC dataset [7, 10]. The sources differ in staining, imaging conditions, annotation detail, and acquisition structure and include both conventional single-cell crops and multi-focus stacks.

The complete curated corpus is publicly available through Zenodo [11]. Source annotations were harmonized into the following 14-class taxonomy: Basophil, Blast, Eosinophil, Erythroblast, Immature WBC, Lymphocyte, Lymphocyte (atypical), Metamyelocyte, Monocyte, Myeloblast, Myelocyte, Neutrophil, Neutrophil (band), and Promyelocyte.

### Study design and operational definitions

We reconstructed analysis groups from relationships recoverable in the available source structure and filenames. In the multi-focus dataset [10], files sharing a numeric cell identifier formed a verified focal stack of ten planes from one physical cell. In Raabin-WBC [3], the final numeric filename suffix was removed and files sharing the remaining base were grouped within class. These groups may reflect shared acquisition context but do not necessarily correspond to individual physical cells, and some may therefore contain distinct cells. Identical filename bases occurring under different class labels were kept separate, while images not covered by either grouping rule were treated as singletons.

Throughout the manuscript, the intact group reference denotes a partitioning condition in which the analysis groups defined above are kept within a single partition. It does not imply independence by patient, smear, specimen, or acquisition field. Random file-level assignment denotes assignment of individual files without preserving these analysis groups. We use performance inflation operationally to describe the difference in apparent test performance between random file-level assignment and the intact group reference under this specific protocol. This term does not imply that every group reconstructed from filenames represents the same physical cell or that the observed difference is a causal effect at the patient level.

### Controlled split generation

For each class, we identified analysis groups containing more than one image. For a dissolution fraction *f* ∈ {0, 0.1, 0.25, 0.5, 1.0}, a deterministic random stream selected *round*(*fm*) of the *m* multi-image groups in that class for dissolution. Images from a dissolved group were then treated as individual assignment units, whereas all remaining groups stayed intact. Thus, *f* = 0 preserved all reconstructed groups and *f* = 1 made every file eligible for independent assignment.

A greedy allocation procedure applied within each class assigned the resulting units to training, validation, and test partitions with target proportions of 70/15/15. Five seeds (1–5) were generated for every dissolution fraction. A separate deterministic random stream was used for each fraction and class, so conditions are independent realizations rather than nested or paired versions of one partition. For every split, we recorded the number and proportion of analysis groups spanning partitions; directional exposure between training, validation, and test sets; and the mean number of training-set group-mates per test image.

### Model training and evaluation

The primary classifier was a torchvision ViT-B/16 initialized with ViT_B_16_Weights.IMAGENET1K_V1. The original classification head was removed. The resulting feature vector was passed through dropout of 0.1, a 128-dimensional linear projection, and a 14-output linear classification layer. The architecture, input resolution, loss, optimization settings, augmentation pipeline, and training schedule were fixed across all ViT partitioning conditions.

Training images were resized to 257 pixels, randomly cropped to 224 × 224 pixels using a scale range of 0.08–1.0 and aspect-ratio range of 0.75–1.33, and horizontally flipped with probability 0.5. Evaluation images were resized to 257 pixels and center-cropped to 224 × 224 pixels. Bilinear interpolation with antialiasing was used. Images were normalized with ImageNet channel means (0.485, 0.456, 0.406) and standard deviations (0.229, 0.224, 0.225).

Models were optimized with AdamW (*β*_1_ = 0.9, *β*_2_ = 0.999, *ɛ* = 10*^−^*^8^). The learning rate was 2 × 10*^−^*^5^ for the pretrained backbone and 10*^−^*^3^ for the projection and classification layers. Weight decay was 0.05 for the backbone and 10*^−^*^4^ for the head. The batch size was 64, and the global gradient norm was clipped at 5.0. We used cross entropy with class weights proportional to *n^−^*^1^*^/^*^2^, normalized to a mean weight of 1, and label smoothing of 0.05. A two-epoch linear warmup was followed by cosine decay to 0.02 of the initial learning-rate multiplier. Training continued for at most 30 epochs and stopped after seven consecutive epochs without improvement in validation macro F1. The checkpoint with the highest validation macro F1 was used for test evaluation.

Accuracy and macro F1 were calculated on a fixed 14-class axis. Macro F1 was the unweighted mean of per-class F1 values. To assess whether the endpoint pattern depended on the primary architecture, we additionally trained a torchvision ResNet-50 initialized with ResNet50_Weights.IMAGENET1K_V2 and a 14-output linear head. ResNet-50 was evaluated for the *f* = 0 and *f* = 1 endpoints using the same split manifests and training protocol, with five seeds per condition.

### Statistical analysis

For each ViT partitioning condition, we report the mean and sample standard deviation across five complete split-and-training runs. Endpoint performance inflation was defined as the difference between the mean random file-level condition (*f* = 1) and the mean intact group reference (*f* = 0). Because the fractions were generated from separate random streams, endpoint conditions were treated as unpaired. We report 95% confidence intervals for the endpoint mean differences using Welch’s unequal-variance *t* procedure. These intervals summarize run-to-run computational and partition variability under the implemented protocol; they do not quantify biological sampling uncertainty, and the five seeds are not biological replicates.

All model training and analysis were run with Python 3.13.7, PyTorch 2.6.0+cu118, torchvision 0.21.0+cu118, NumPy 2.1.2, and Pillow 11.0.0.

### Descriptive exposure analysis under random file-level assignment

To provide additional descriptive context for the endpoint comparison, we trained five additional ViT-B/16 models under random file-level assignment (*f* = 1), one for each seed, using the same split manifests and training recipe as the primary ViT endpoint. Within each run, a test image was defined as *exposed* when at least one image from the same analysis group occurred in the training partition; all other test images were *unexposed*. Both subsets were evaluated with the corresponding model.

Because exposure was associated with source, class, and group type, exposed and unexposed subsets were not exchangeable. We therefore treat all exposure comparisons as descriptive rather than causal. We report accuracy and macro F1 on the fixed 14-class axis. For the within-class comparison, an accuracy difference was calculated only when a class contained at least 20 exposed and 20 unexposed test images in a given seed; eligible class differences were then averaged within that seed.

### Data and code availability

The curated image corpus is available on Zenodo at https://doi.org/10.5281/zenodo.19126698 [11]. The HSD-Bench software and reproducibility archive are available at https://doi.org/10.5281/zenodo.22807823 with the development repository at https://github.com/lowoncuties/HSD-Bench [12]. The archive contains the pinned experiment definition, deterministic split-generation code, model-training implementation, integrity checks, exact partition digests, grouping summaries, run-level overall metrics, per-class summaries, confusion-matrix summaries, relatedness summaries, exposure-analysis outputs, and the source data used to generate the figures retained for the reported study. Original image data remain subject to the terms of their upstream source datasets.

### Ethics statement

No new participants or biological specimens were recruited or collected for this study. The analyses used publicly available image datasets under their stated access conditions, and the authors did not receive or analyze identifiable personal information. The present work is a secondary computational analysis of those resources; no additional intervention or participant contact was performed.

## Results

The corpus comprised 42,362 reconstructed analysis groups. Of these, 11,095 (26.2%) contained more than one image and together accounted for 97,906 images (75.8% of the corpus). These groups included 9,419 verified ten-plane focal stacks and 1,676 Raabin groups reconstructed from filenames. Verified focal-stack members are repeated views of one physical cell, whereas Raabin groups reconstructed from filenames may reflect a shared acquisition context and can include distinct cells. The remaining 31,267 groups were singletons.

### Controlled partitions produce a graded range of relatedness

Across the five partitioning conditions, realized relatedness increased systematically while the same source corpus and class-stratified 70/15/15 target proportions were used (Table 1). No reconstructed analysis group crossed a partition boundary in the intact group reference. Under random file-level assignment, 75.1% of test images had at least one training-set group-mate. Intermediate conditions produced test exposure levels of 16.0%, 40.2%, and 72.9%.

**Table 1.** Controlled split conditions (five seeds each) and mean realized relatedness. Arrows denote the partition containing a group-mate; for example, Val ← Train is the percentage of validation images with a training-set group-mate.

| Nominal $f$ | Groups spanning | Val ← Train | Test ← Train | Test ← Val | Mean training group-mates per test image |
| --- | --- | --- | --- | --- | --- |
| 0.0 | 0.0% | 0.0% | 0.0% | 0.0% | 0.00 |
| 0.1 | 2.4% | 15.9% | 16.0% | 15.8% | 0.49 |
| 0.25 | 6.1% | 40.0% | 40.2% | 39.6% | 1.22 |
| 0.5 | 12.2% | 73.0% | 72.9% | 69.8% | 2.64 |
| 1.0 | 23.5% | 75.0% | 75.1% | 56.7% | 4.62 |

Relatedness in the validation set increased together with relatedness in the test set. Because validation macro F1 determined checkpoint selection, the contrasts below quantify the complete partitioning protocol rather than an isolated causal effect of training–test relatedness alone.

### Random file-level assignment yields higher apparent classifier performance

For ViT-B/16, random file-level assignment increased mean test macro F1 by 0.124, from 0.853 to 0.977, and mean test accuracy by 0.139, from 0.851 to 0.990, relative to the intact group reference (Table 2). Under the operational definition used in this study, these differences represent performance inflation associated with relaxing the reconstructed grouping constraints.

**Table 2.** ViT-B/16 performance under the intact group reference and random file-level assignment (five independent split-and-training runs per condition; mean ± sample standard deviation). The confidence intervals summarize computational and partition variability across runs and are not biological-sampling intervals.

| Split | Test macro F1 | Test accuracy |
| --- | --- | --- |
| intact group reference ($f = 0$) | 0.853 ± 0.010 | 0.851 ± 0.008 |
| Random file-level assignment ($f = 1$) | 0.977 ± 0.003 | 0.990 ± 0.004 |
| **$\Delta$, 95% confidence interval** | **+0.124 (0.112–0.137)** | **+0.139 (0.129–0.149)** |

Apparent performance increased progressively across the intermediate conditions. Mean test macro F1 was 0.853, 0.878, 0.922, 0.965, and 0.977 for *f* = 0, 0.1, 0.25, 0.5, and 1.0, respectively. The corresponding realized proportions of test images with at least one training-set group-mate were 0%, 16.0%, 40.2%, 72.9%, and 75.1%. Although binary exposure changed little between *f* = 0.5 and *f* = 1.0, the mean number of training group-mates per test image increased from 2.64 to 4.62. Thus, the endpoint difference was accompanied by a graded relationship across the partition family rather than arising only from a comparison of two endpoints (Fig 2).

**Fig 2.**
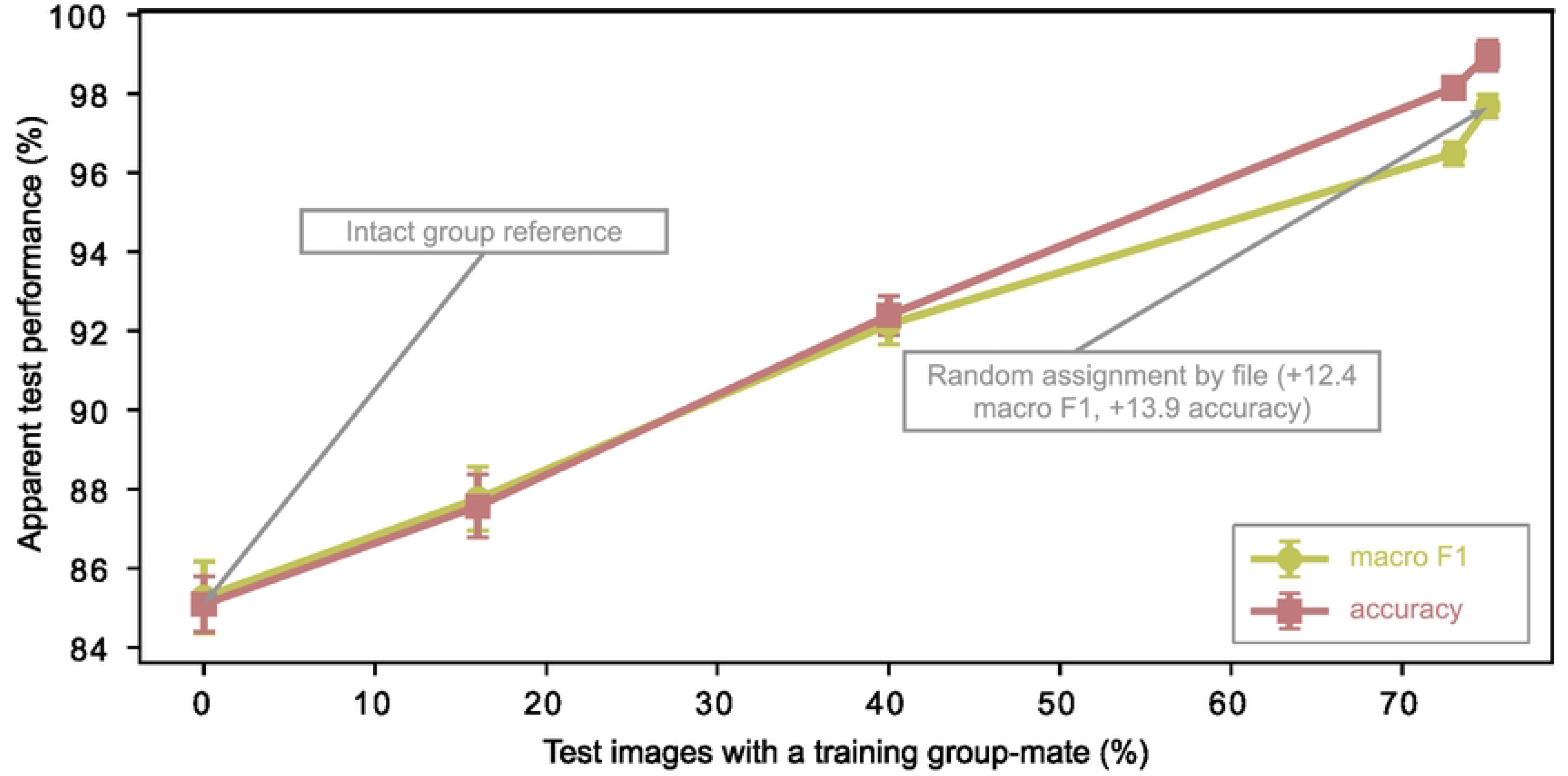
Graded relationship between realized train–test relatedness and apparent benchmark performance. Points show the mean of five independent split-and-training runs per condition, and error bars show sample standard deviation. The *x*-axis is the realized fraction of test images with at least one member of the same analysis group in the training set. Macro F1 and accuracy increase across the controlled partitioning conditions. Because validation relatedness changes concurrently with test relatedness, the figure describes sensitivity to the full partitioning protocol and should not be interpreted as an isolated causal dose–response effect.

### The endpoint pattern appears with two model families

The same endpoint pattern was observed with the convolutional ResNet-50 backbone (Table 3). Random file-level assignment increased ResNet-50 macro F1 by 0.119 and accuracy by 0.130, compared with increases of 0.124 and 0.139 for ViT-B/16. The direction and magnitude of the difference associated with partitioning were therefore similar for the two evaluated model families.

**Table 3.** Endpoint performance for transformer and convolutional backbones (five independent split-and-training runs per condition; mean ± sample standard deviation).

| Model | Split | Macro F1 | Accuracy |
| --- | --- | --- | --- |
| ViT-B/16 | $f = 0$ | $0.853 \pm 0.010$ | $0.851 \pm 0.008$ |
| ViT-B/16 | $f = 1$ | $0.977 \pm 0.003$ | $0.990 \pm 0.004$ |
| ResNet-50 | $f = 0$ | $0.848 \pm 0.007$ | $0.847 \pm 0.008$ |
| ResNet-50 | $f = 1$ | $0.967 \pm 0.002$ | $0.977 \pm 0.001$ |
| **Inflation ($\Delta$, ViT-B/16)** |  | **+0.124** | **+0.139** |
| **Inflation ($\Delta$, ResNet-50)** |  | **+0.119** | **+0.130** |

### The largest class-level differences occur in immature and blast classes

The endpoint difference was not uniform across cell types. The largest changes occurred for Promyelocyte, whose mean F1 increased from 0.488 to 0.899 (ΔF1 = +0.411); Metamyelocyte, from 0.615 to 0.985 (ΔF1 = +0.369); and Blast, from 0.627 to 0.990 (ΔF1 = +0.363). In contrast, the differences were +0.004 for Neutrophil, +0.010 for Lymphocyte, and +0.015 for Eosinophil (Table 4). We do not use the correlation between intact group F1 and ΔF1 as independent evidence because ΔF1 mathematically contains the negative of the intact group F1 term.

**Table 4.**
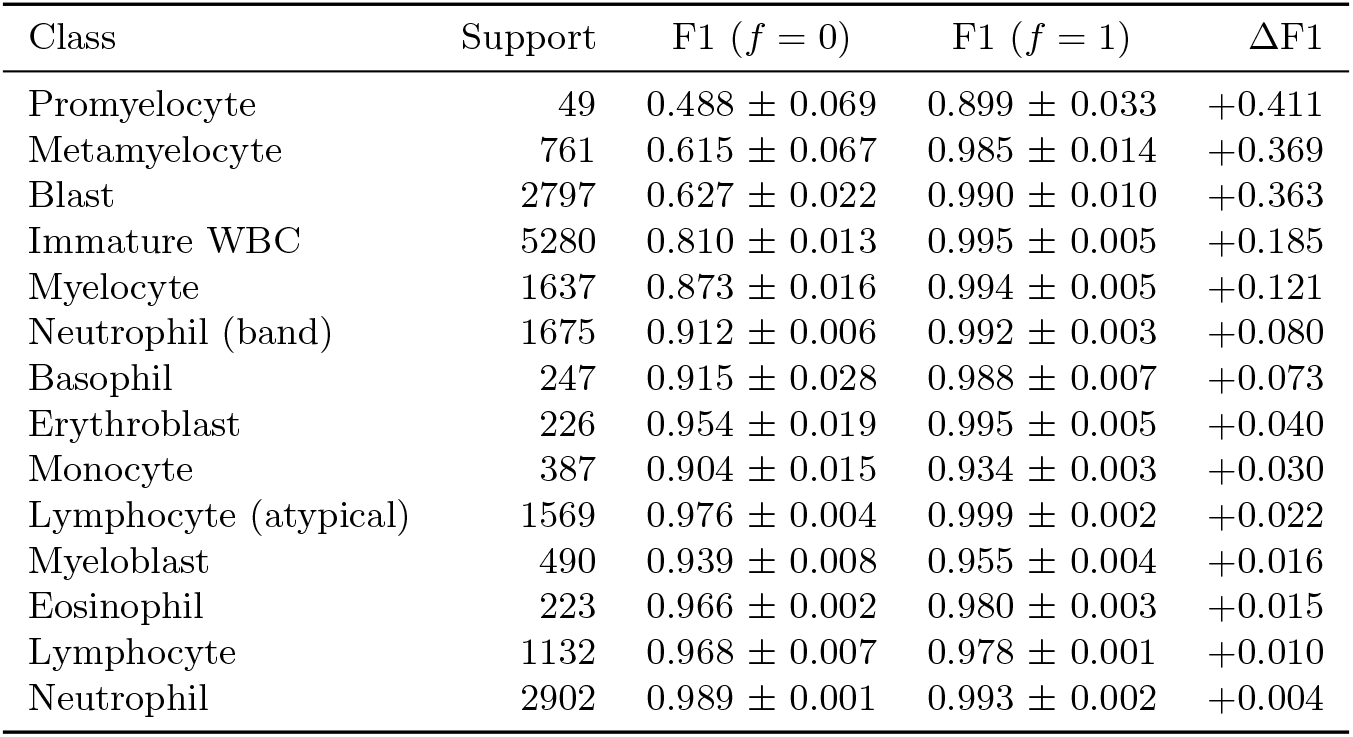
Per-class ViT-B/16 test F1 under the intact group reference (*f* = 0) and random file-level assignment (*f* = 1), sorted by ΔF1. Values are mean ± sample standard deviation over five independent split-and-training runs. Support is the mean number of test images under the intact group reference.

The row-normalized confusion matrices show the same qualitative pattern (Fig 3). Confusions among several immature granulocytic and blast-related classes that are visible under the intact group reference are markedly reduced under random file-level assignment.

**Fig 3.**
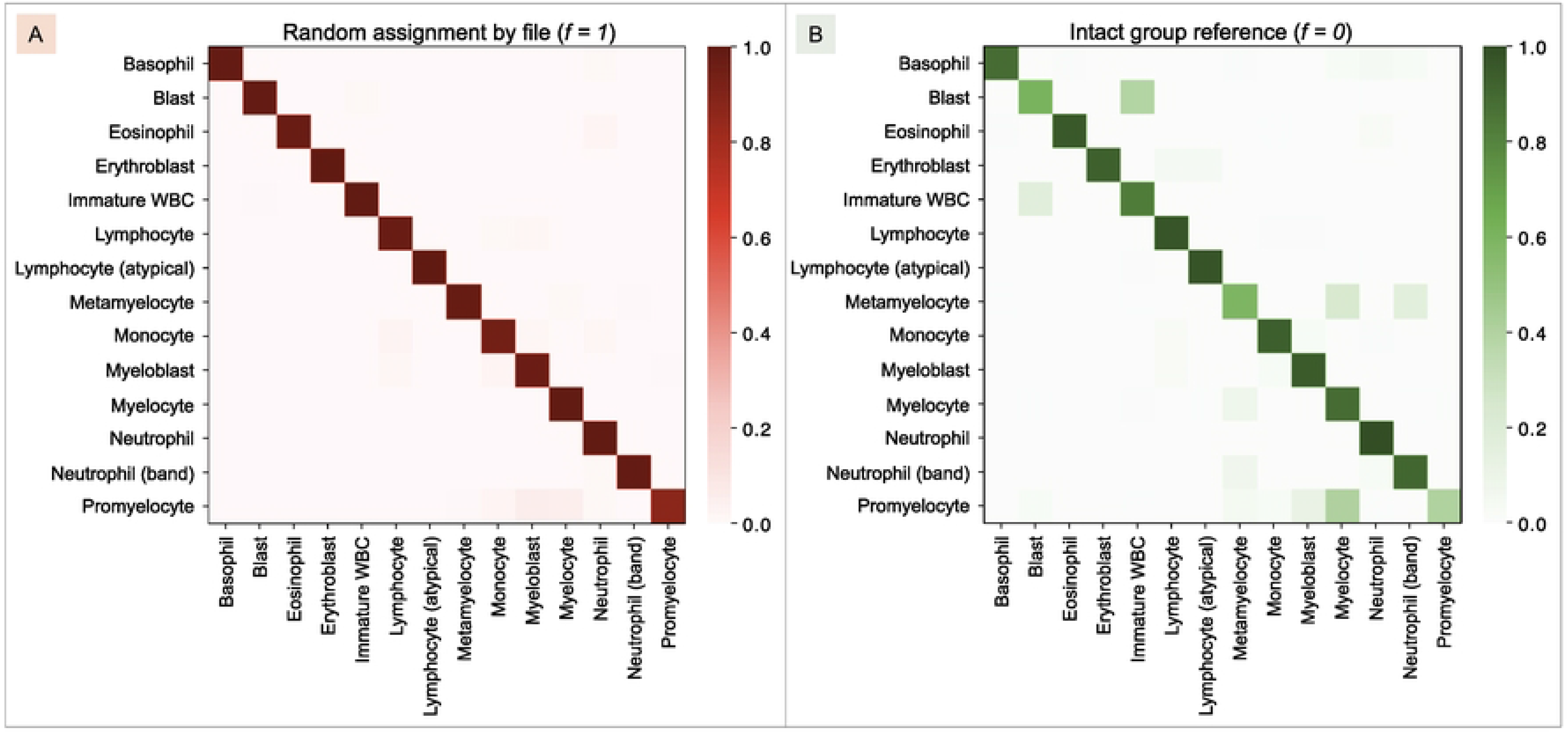
Confusion matrices under the intact group reference and random file-level assignment. Row-normalized confusion matrices are averaged over five ViT-B/16 runs per condition. Rows denote true classes and columns denote predicted classes. The color scale ranges from 0 to 1 after row normalization. The intact group reference shows additional off-diagonal confusion among several immature granulocytic and blast-related classes, whereas these confusions are reduced under random file-level assignment.

### Descriptive within-split exposure analysis

Within the five additional random-assignment runs, exposed and unexposed test subsets differed substantially in composition. Exposed images were enriched for multi-image groups and classes containing many focal stacks, whereas unexposed images contained a larger proportion of classes represented mainly by singleton groups. Raw between-subset comparisons therefore do not identify a counterfactual effect of exposure.

Across seeds, the mean exposed-minus-unexposed accuracy difference after averaging within eligible classes was +0.114, with seed-level values ranging from +0.013 to +0.203; four or five classes met the support threshold in each seed (Fig 4A). On the fixed 14-class axis, mean macro F1 was 0.907 for exposed images and 0.618 for unexposed images, compared with 0.853 for the intact group reference (Fig 4B). Mean raw accuracy was 0.997 for exposed images and 0.976 for unexposed images. Because exposure covaried with source, class, and group type, these values are descriptive and should not be interpreted as a causal estimate.

**Fig 4.**
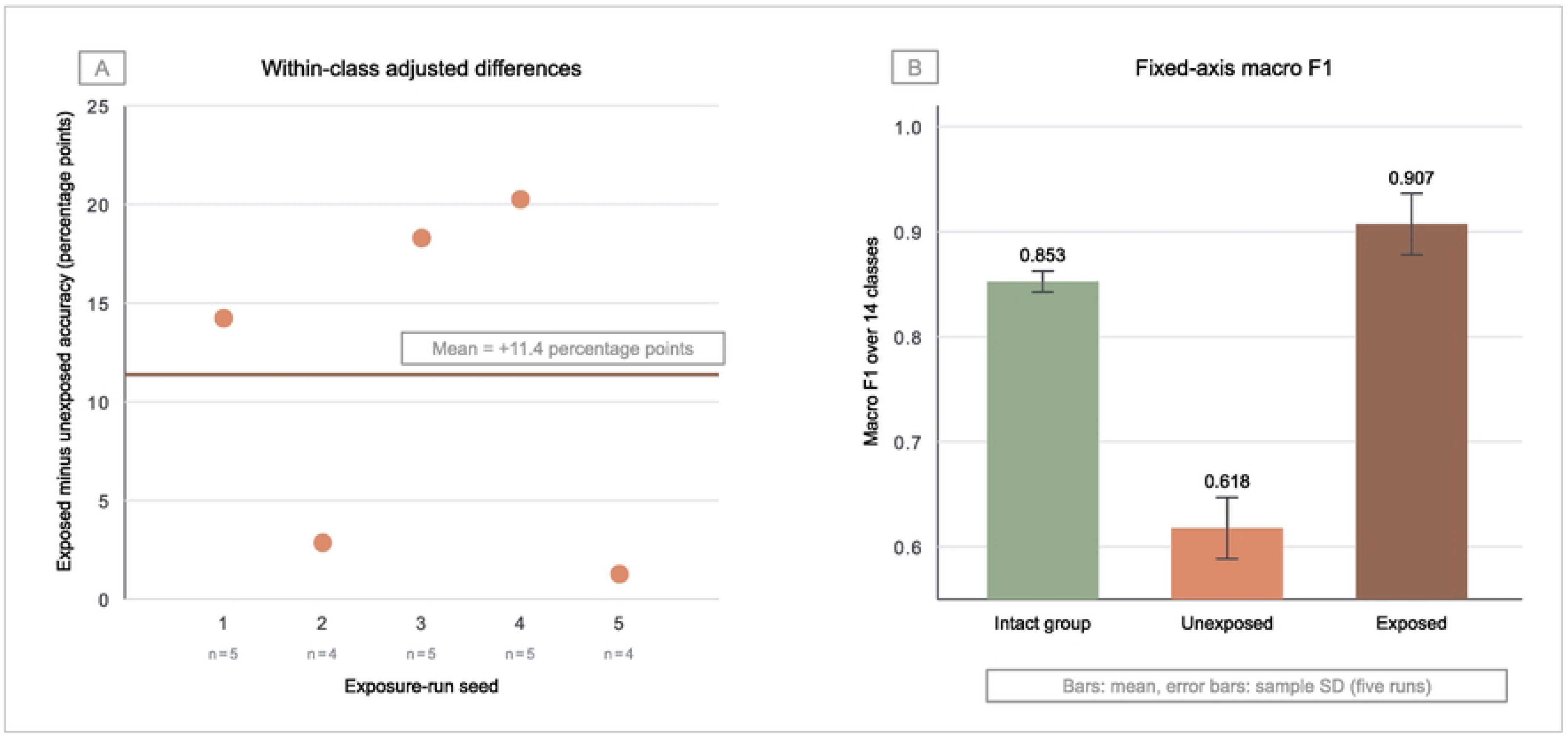
Descriptive exposure analysis under random file-level assignment (*f* = 1). Five separately trained ViT-B/16 runs are summarized, with the intact group reference shown for context. (A) For each seed, the plotted value is exposed minus unexposed accuracy averaged across classes containing at least 20 images in both subsets. The horizontal line marks the five-run mean of 0.114 (11.4 percentage points); four or five classes qualified per seed. (B) Macro F1 is calculated on the same fixed 14-class axis for the intact group test set and the unmatched exposed and unexposed subsets. Bars show means and error bars show sample standard deviations across five runs. The exposed and unexposed subsets differ in source, class, and group type, so this analysis is descriptive and does not identify a causal effect.

## Discussion

This study measured how classification performance changes when related samples are increasingly allowed to appear in different partitions of a biomedical image benchmark. The benchmark combined images from multiple sources. When all identified groups were kept intact, ViT-B/16 achieved a mean macro F1 of 0.853 and an accuracy of 0.851. With random assignment of individual files, these values increased to 0.977 and 0.990. Performance increased gradually as fewer groups were kept intact. ResNet-50 showed a similar difference between the two conditions. The largest differences occurred in several immature granulocytic and blast classes.

These findings extend previous work on data leakage and evaluation with dependent samples [1, 2]. Earlier studies have shown that shared information between training and evaluation data can inflate performance estimates. Here, we examined how these estimates change as progressively fewer sample groups are preserved during partitioning. The source images and training procedure remained the same across conditions. The results show that measured performance can increase gradually as more related samples appear across partitions. They also show that overall scores can hide substantial differences between classes.

The results for individual classes are relevant because the benchmark is imbalanced. Several of the largest increases occurred in immature or blast categories that were difficult to classify when groups were kept intact. If a class contains few independent groups, placing related observations in both training and evaluation data can make it appear much easier to classify. Mature classes that were already well separated showed little change. The descriptive exposure analysis supports this interpretation, but the exposed and unexposed subsets also differed in source, class, and group type. This comparison therefore cannot show how the same test image would have been classified if related observations had been excluded from training.

The appropriate unit for splitting data depends on the task. If the aim is to classify previously unseen cells, all focal planes and derived images of the same physical cell should remain in one partition. Separating patients imposes a stricter requirement. This is particularly relevant when predicting patient outcomes or when acquisition characteristics associated with individual patients could help the model make predictions. Separation by smear, specimen, slide, or acquisition may also be needed when the study aims to generalize across those units. A numerical split ratio alone does not explain which of these forms of separation was used.

Studies in blood cell imaging illustrate both the risks and the available safeguards. Matek et al. [13] used cross-validation based on individual images and screened for overlapping image regions. Their approach illustrates why splitting individual images does not, by itself, establish that the same cell appears in multiple partitions. Park et al. [7] provide metadata linking images taken at different focal planes. These metadata allow all images of one cell to remain together and support stricter separation by slide or patient when needed. Raabin-WBC provides coordinates, acquisition metadata, and procedures for handling duplicate observations [3]. Other recent systems evaluate performance on external datasets or separate data by patient [14–16]. These examples support reporting which biological or acquisition unit was kept separate, alongside the split ratio.

Dataset preparation can make relationships between samples harder to identify. Augmentation, object extraction, repeated cropping, and redistribution through secondary repositories can obscure which files share a common source. Random image splitting alone is not evidence that a published study contains leakage. The studies discussed here illustrate reporting practices and safeguards, rather than establish leakage in individual published results. Reports should state whether data were split before or after augmentation and object extraction. They should also explain whether images derived from the same source remained together and what type of generalization the evaluation supports.

HSD-Bench provides a reproducible way to measure this sensitivity in the present corpus. Its grouping metadata allow researchers to generate both a reference split that keeps all identified groups intact and alternative splits that preserve fewer groups. The released manifests document the exact assignment of samples to partitions. The same approach could be applied to datasets containing repeated measurements, multiple views, patches, or other derived samples. However, the rules for identifying related samples depend on the dataset and domain. Applying the workflow to another dataset therefore requires defining and validating the grouping unit.

For biomedical image benchmarks, we recommend reporting the unit used to separate training, validation, and test data, the identifiers or metadata used to link related observations, and the level of generalization that the split supports. External validation, evaluation on separate patients, and predefined split manifests can strengthen a study. Each internal evaluation should still state clearly which biological or acquisition units were kept separate.

## Limitations

The quantitative experiment evaluates one curated blood cell corpus. The magnitude of the observed differences should therefore not be generalized numerically to other datasets, tasks, acquisition protocols, or class distributions. Validation-set relatedness also changes with test-set relatedness because each condition defines a complete train/validation/test protocol and validation macro F1 is used for checkpoint selection. The experiment consequently estimates sensitivity to the full partitioning protocol rather than an isolated causal effect of training–test exposure. The five seeds quantify computational and partition variability and are not biological replicates.

## Conclusions

In this biomedical image classification benchmark, apparent test performance increased progressively as members of recoverable sample groups were increasingly allowed to occur across dataset partitions. The largest apparent gains were concentrated in several difficult immature granulocytic and blast classes, and the same overall pattern was observed with both transformer and convolutional backbones. These findings show that the unit of independence should be considered an explicit part of benchmark design rather than a secondary implementation choice. Clear grouping metadata, transparent partition definitions, and explicit reporting of the intended unit of generalization can improve the interpretation, comparability, and reproducibility of biomedical machine learning results.

## Acknowledgments

The authors have no acknowledgments to report.

## Notes

### Competing Interest Statement

The authors have declared no competing interest.

